# Plasma glutathione status as indicator of pre-analytical centrifugation delay

**DOI:** 10.1101/2020.12.09.417386

**Authors:** Tamara Tomin, Natalie Bordag, Elmar Zügner, Abdullah Al-Baghdadi, Maximillian Schinagl, Ruth Birner-Gruenberger, Matthias Schittmayer

## Abstract

Prolonged incubation of blood prior to plasma preparation can significantly influence the quality of the resulting data. Different markers for this pre-clinical variability have been proposed over the years but with limited success.

In this study we explored the usefulness of glutathione (GSH) status, namely ratio of reduced to oxidized glutathione (GSH/GSSG), as potential marker of plasma preparation delay. For that purpose, blood from 20 healthy volunteers was collected into tubes with a cysteine quencher (N-ethylmaleimide; NEM) for GSH stabilization. Plasma preparation was delayed at room temperature for up to 3 hours and every hour, a plasma sample was prepared and the GSH/GSSG ratio measured.

We report that over the course of the investigation, plasma concentrations of both GSH and GSSG increased linearly (R^2^ = 0.99 and 0.98, respectively). Since GSH increased at a much faster rate compared to GSSG, the GSH/GSSG ratio also increased linearly in a time dependent manner (R^2^ = 0.99). As GSH is an intracellular antioxidant, we speculated that this might stem from ongoing blood hemolysis, which was confirmed by the time dependent rise in lactate dehydrogenase (LDH) activity in the plasma samples. Moreover, we demonstrate that the addition of the thiol alkylating reagent NEM directly to the blood tubes does not seem to influence downstream analysis of clinical parameters. In conclusion we propose that the glutathione status could be used as an indicator of the centrifugation delay prior to plasma preparation.

## INTRODUCTION

In order to minimize pre-analytical variability, standard operating procedures (SOPs) for plasma and serum preparation have been continuously re-evaluated and discussed over the last decades (*1*). Reasons for concern are that several factors involved in pre-analytical processing can be the critical determinants of the quality of obtained results. One such prominent variable is the pre-centrifugation delay, i.e. the lag time between blood collection and plasma preparation (*2*), which is shown to greatly influence outcome of the various analysis. According to a broad metabolite profiling study conducted by Kamlage et *al*. (*3*), a delay of 2 h at room temperature (RT) prior to plasma preparation affected 22 % of all monitored metabolites. In comparison, inducing hemolysis of grade 1 (passing the blood through a 25-gauge needle) affected only 18 % of all metabolites included in the study (*3*). Prominent changes in the metabolome were also reported by Bernini et *al*. (*4*), who demonstrated that the plasma metabolic profile was highly dependent on the time between blood collection and plasma preparation. Besides impacting the metabolome, delays in plasma (or serum) preparation also affect the proteome. Both serum and plasma proteome profiles are significantly altered by processing delays (*5*). Hsieh et *al*. showed that prolonged delays at RT led to significant increase in the variation (coefficient of variation (CV) > 20 %) of 66 % of all analyzed peaks (*5*).

To better assess the effects of prolonged incubation time between blood collection and plasma preparation, efforts have been made to discover potential indicators of blood processing delay. Lee et *al*. (2015) observed that if blood separation was postponed by 48 h, plasma content of inorganic phosphate and potassium rose prominently (*6*). Similarly, the same group also suggested a panel of cytokines (L-1β, GM-CSF, sCD40L, IL-8, MIP-1α, and MIP-1β) as potential indicators of the delay in plasma and serum preparations (*7*). A comprehensive review of proposed preparation delay markers was published by Ruiz-Godoy et *al*. (2019) (*2*).

Nevertheless, there is a lack for a standardized, sensitive, stand-alone marker delivering a clear read-out of the delay in plasma processing. Such a marker would have to be stable and would need to be detectable in plasma already after one or two hours processing wait, as those time points were shown to already significantly change analysis results (*3, 4*). In an attempt to tackle this conundrum, in the present study we focused on the influence of plasma preparation delay on small molecular thiols, namely the ratio of reduced to oxidized glutathione. Glutathione (GSH) is one of the most abundant intracellular thiols and the ratio of GSH to oxidized glutathione (glutathione disulfide, GSSG) is often used as a readout of tissue oxidative state (*8, 9*). Addressing GSH and GSSG in biological fluids represents a challenge for itself, as GSH is prone to artificial oxidation during sample preparation (*9, 10*). However, if properly stabilized with a thiol-blocking (alkylating) agent at the point of sample collection, precise quantification can be ensured (*9–11*).

While treatment with alkylating reagents stabilizes the intracellular glutathione (*9, 10*), the aim of this study was to investigate how a delay in blood separation of alkylated blood can influence glutathione status of extracellular fluids, in particular plasma. To achieve this, we spiked blood collection tubes with N-ethylmaleimide (NEM), a potent thiol-blocking reagent (*11, 12*), and incubated the blood for up to three hours at room temperature prior centrifugation and plasma collection. Every hour (60 min) an aliquot of blood was taken and spun down to collect plasma, which was then used to monitor the GSH/GSSG ratio over time.

## MATERIALS AND METHODS

### Study design

The study was conducted in adherence to the Declaration of Helsinki and was reviewed by the ethical committee of the Medical University of Graz, Austria (31-116 ex 18/19, 16.01.2019). Prior to commencement of any study activities a written informed consent from all participants was obtained.

A study-subgroup of 20 healthy volunteers (10 female, 10 male) was carefully selected to reduce biological and metabolic variation. Inclusion criteria were healthy Caucasian male or female, age from 20 to 30 years (mean 26 ± 3 S.D.), body mass index (BMI) from 18.5 to 25 kg/m^2^ (mean 22 ± 2 S.D.), abstinent of drug abuse for >1 year pre-trial, nonsmoker or light smoker (≤1 cigarette/week), abstinent or light alcohol consumption (≤7 units/week, 1 unit = 10 ml or 8 g alcohol), overnight fasting (12h), and training abstinence (24h). Exclusion criteria included any acute, or chronic diseases, hormonal contraception, medication with heparin (nonsteroidal or steroidal anti-inflammatory) in the last ten days, medication with antihistamines or selective serotonin reuptake inhibitors in the last four weeks, *in vitro* fertilization (IVF) treatment or any surgery in the last three months, any special diet form (e.g. vegan, gluten-free, malabsorption specific diets, ketogenic…) or any other condition which would interfere with the safety of the participant, especially known anemia, blood or plasma donation within the last month, pregnancy, breastfeeding, intention of becoming pregnant or not using adequate contraception, mental incapacity, unwillingness or language barriers precluding adequate understanding or co-operation. All samples were collected within 3 weeks. Detailed volunteers’ characteristics can be found in Suppl. Data 1.

### Sample collection and plasma preparation

Blood was collected in the morning between 08-10 a.m. (MESZ) with a 21G butterfly needle while sitting. Venipuncture was performed maximum once per arm. The tourniquet was released after 1 min and blood collection was achieved in 4 to 9 min. Several tubes were collected (e.g. for clinical routine hematology or biobank storage); the first tube was discarded and each tube was immediately inverted gently 3 times.

### NEM stabilized sample collection and preparation

Shortly before the blood collection, VACUETTE^®^ (Grainer Bio-One, AT) 3 ml K_3_EDTA blood collection tubes were spiked with 100 µl of 75 mM N-ethylmaleimide (NEM) in phosphate saline buffer (PBS) (final NEM concentration 2.5 mM). After blood collection, tubes were kept upright at room temperature and at regular intervals (0, 60, 120 and 180 min) 500 µl of NEM stabilized blood was transferred from the blood collection container to a 1.5 ml microcentrifuge tube using a 1 ml trimmed pipet tip to minimize cell damage. Microcentrifuge tubes were then spun down for 10 min at 1300 x *g* according to standard operating procedure (SOP) (*1*) for plasma preparation and 100 µl of plasma was carefully transferred to a new microcentrifuge tube and frozen till further analysis.

To analyze whether NEM affects common blood parameters, blood from 20 individuals was collected in VACUETTE^®^ (Grainer Bio-One, Austria) 5 ml Lithium Heparin Tubes spiked with either 100 µl of 200 mM NEM in PBS (2.5 mM final concentration) or 100 µl PBS as vehicle control shortly before sample collection.

### Clinical routine hematology

Tubes for clinical routine hematology were transported (10-15 min) in isolated boxes, the temperature was logged and did not exceed the limits of 20-25°C. The transport was within 2.5 h after phlebotomy and hematology measurement was performed within 4 h at the Clinical Institute of Medical and Chemical Laboratory Diagnostics, Medical University of Graz. Tubes from the same volunteer were always handled equally during transport and subsequent measurement. Hematological measurements were carried out on Sysmex Blood counters of XN or XE series (Sysmex Co., Japan) through means of flow cytometry. All molecular analyses were carried out on a COBAS 8000 instrument (Roche Diagnostics, Switzerland) with following methodologies applied: ions were measured through indirect potentiometry; alanine and aspartate aminotransferase, as well as glucose, creatinine and cholesterol were addressed spectrophotometrically; troponin T and thyroid-stimulating hormone via electro-chemiluminescence immunoassay (ECLIA) and C-reactive protein with an immunological turbidity test.

### Glutathione measurements

Reduced (GSH) and oxidized glutathione (glutathione disulfide, GSSG) were addressed using a two-step alkylation protocol followed by liquid-chromatography coupled to mass spectrometry analysis (LC-MS/MS) as previously described (*9*) with a difference that the measurement was carried out on a TSQ Access Max triple quadrupole (Thermo Scientific, USA) operating in positive SRM mode. TSQ parameters were: spray voltage 3000 V, capillary temperature 240 °C, vaporizer temperature 300 °C, and sheath gas; ion sweep gas and aux gas pressures were 35, 0 and 5 units, respectively. Tube lens offset was set to 48 for the lower molar masses and skimmer offset was set to 0. The transitions with their corresponding collision energies are listed in Table 1.

**Table 1.** Multiple reaction monitoring (MRM) table of transitions for glutathione and glutathione derivatives. Transitions highlighted in bold and italic were used for quantitation.

| Metabolite | Parent ion | Product ion | Collision energy |
| --- | --- | --- | --- |
| ***GSH*** | ***308*** | ***179*** | ***19*** |
| ***GSH-NEM (Transition 1)*** | ***433.16*** | ***201*** | ***22*** |
| GSH-NEM (Transition 2) | 433.16 | 304 | 15 |
| ***GSH-d5-NEM (Transition 1)*** | ***438.16*** | ***206*** | ***22*** |
| GSH-d5-NEM (Transition 2) | 438.16 | 309 | 15 |
| ***<sup>13</sup>C<sub>2</sub>, <sup>15</sup>N-GSH-d5-NEM (IS)*** | ***441.16*** | ***206*** | ***22*** |
| ***GSSG*** | ***613.2*** | ***355.3*** | ***10*** |

### Lactate dehydrogenase (LDH) activity assay

LDH activity of 14 plasma samples prepared after the indicated time delay (0, 60, 120 or 180 min after the blood collection) was addressed spectrophotometrically using Lactate Dehydrogenase Activity Assay Kit (MAK066, Sigma Aldrich, USA) according to the manufacturer’s instructions. Briefly, per each well 25 µl of plasma was mixed with 25 µl of assay buffer and added to 50 µl of reaction mix (consisted of substrate mix diluted in the assay buffer). On each microtiter plate duplicates of 3 µl of positive control were included. After the initial read, the plate was read continuously every five minutes for the total length of 20 min. Absorption values obtained after 10 min of incubation (per each time point) were all in the linear range and selected for quantification.

### Statistical analysis

Data visualization and statistical analysis were performed with Microsoft Excel and R (*13*) (v4.0.2, packages plyr, stringr, readxl, ggplot2, dendsort, pheatmap, cellWise, Metaboanalyst R) (*14, 15*). Statistical significance of GSH/GSSG or LDH activity was tested by applying one-way ANOVA or Student t-tests with a p-value of 0.05 as the significance threshold. If not stated otherwise, values are displayed as mean values ± standard error of mean (S.E.M).

For the analysis of NEM influence on clinical parameters data distribution and scedasticity were investigated with Kolmogorov-Smirnov test and Brown-Forsythe Levene-type test, respectively, and multiple test adjustment by Benjamini-Hochberg (BH) (Suppl. Data 1). From the 26 parameters three had to be excluded because the data behaved categorical (Baso, Eos, Mono, for abbreviations see Table 2) and two had to be excluded due to too many missing values (below detection limit, CRP, TNTHS). All remaining 21 numeric parameters were found to be sufficiently normal distributed and homoscedastic for further statistical analysis without transformation.

**Table 2.** Detailed list of blood parameters from blood with PBS or NEM addition. The mean was calculated pairwise (one mean value per volunteer (n=20) from both tubes). CV_w_ represents the mean of all standard deviations obtained per volunteer divided by the mean value of the given blood parameter. Parameters which were not part of the statistical analysis are highlighted in italic.

| abbrevia<br>tion | full name | mean<br>(PBS&NEM) | CV<sub>w</sub> (%) | reference range |  | unit |
| --- | --- | --- | --- | --- | --- | --- |
|  |  |  |  | male | female |  |
| **Ery** | erythrocytes | 4.63 ± 0.1 | 2.1 | 4.5-5.9 | 4.1-5.1 | 10<sup>12</sup>/L |
| **Leu** | leucocytes | 5.05 ± 0.13 | 2.6 | 4.4-11.3 |  | 10<sup>9</sup>/L |
| **Lym** | lymphocytes | 1.8 ± 0.06 | 3.1 | 1-4.8 |  | 10<sup>9</sup>/L |
| ***Mono*** | *monocytes* | *0.3* |  | *0.2-1.0* |  | *10<sup>9</sup>/L* |
| ***Eos*** | *eosinophils* | *0.2* |  | *0-0.7* |  | *10<sup>9</sup>/L* |
| ***Baso*** | *basophils* | *0.0* |  | *0-0.2* |  | *10<sup>9</sup>/L* |
| **Neutro** | neutrophils | 2.68 ± 0.07 | 2.8 | 1.8-7.7 |  | 10<sup>9</sup>/L |
| **Thromb** | platelets | 203 ± 11 | 5.5 | 140-440 |  | 10<sup>9</sup>/L |
| **ALT** | alanine aminotransferase | 17.3 ± 2.01 | 11.7 | 0-45 | 0-35 | U/L |
| **AST** | aspartate aminotransferase | 18.7 ± 1.06 | 5.7 | 0-35 | 0-30 | U/L |
| **Crea** | creatinine | 0.85 ± 0.02 | 2.3 | 0-1.2 | 0-1 | mg/dL |
| **Cl** | chloride ions | 103 ± 0.39 | 0.4 | 95-110 |  | mmol/L |
| **K** | potassium ions | 4.03 ± 0.11 | 2.6 | 3.5-5.0 |  | mmol/L |
| **Na** | sodium ions | 140 ± 0.25 | 0.2 | 135 | 145 | mmol/L |
| **Chol** | cholesterol | 155 ± 1.7 | 1.1 | 0-200 |  | mg/dL |
| **Glc** | glucose | 92.3 ± 0.78 | 0.8 | 70-100 |  | mg/dL |
| **Hb** | hemoglobin | 13.6 ± 0.28 | 2.1 | 13-17.5 | 12-15.3 | g/dL |
| **HCT** | hematocrit | 39.3 ± 0.7 | 1.8 | 40-50 | 35-45 | % |
| **MCH** | mean cell hemoglobin | 29.3 ± 0.15 | 0.5 | 28-33 |  | pg |
| **MCHC** | mean corpuscular hemoglobin concentration | 34.5 ± 0.29 | 0.5 | 33-36 |  | g/dL |
| **MCV** | mean corpuscular volume | 84.9 ± 0.32 | 0.3 | 80-98 |  | fL |
| **MPV** | mean platelet volume | 10.8 ± 0.74 | 2.9 | 7-13 |  | fL |
| **RDWCV** | Red blood cell distribution width | 12.7 ± 0.24 | 1.9 | 11-16 |  | % |
| **TSHL** | thyroid-stimulating hormone | 2.34 ± 0.03 | 1.1 | 0.10-4.00 |  | μU/ml |

Principal component analysis allowing for missing values and cellwise & rowwise outliers (R function *MacroPCA*) was performed centered and scaled to unit variance (*16*). The number of components was set to cumulatively retain 80 % of explained variance, here six. Orthogonal projections to latent structures discriminant analysis (OPLS-DA) was performed centered and scaled to unit variance (R function *Normalization* with *scaleNorm=“AutoNorm”* and R function *OPLSR*.*Anal*) with a standard 7-fold cross validation for the classification factor *gender* or *NEM*. Model stability was additionally verified with 1000 random label permutations. Detailed model results of MacroPCA and OPLS-DA, including scores, loadings and S plot values, are given in Suppl. Data 1. Hierarchical clustering analysis was performed centred and scaled to unit variance (R function *scale*) per parameter. The dendrograms were clustered by Lance-Williams dissimilarity update with complete linkage (R function *dist* and *hclust*) and sorted (R function *dendsort*) at every merging point according to the average distance of subtrees and plotted at the corresponding heat maps (R function *pheatmap*).

## RESULTS

### Glutathione content in plasma increased with the delay in plasma preparation

Glutathione (GSH) is one of the most abundant intracellular antioxidants, with especially high concentration levels in red blood cells (in the range of 1.2-1.5 mM) (*9, 17*). As the concentration of GSH in the blood cells is up to several folds higher compared to its concentration in physiological fluids such as plasma or serum (*8, 9*), even a slight “leakage” of glutathione from the blood cells can lead to a prominent increase in the GSH content of the liquid blood components. We detected a significant and strong increase of both GSH as well as GSSG in plasma by approximately 3.5 and 1.9 folds, respectively (Fig. 1A), after a delay of plasma preparation at RT for up to 3 h. The increase in GSH content in plasma was continuously higher than the increase in GSSG, so that the GSH/GSSG ratio also rose steadily over time (Fig. 1A). This increase in GSH/GSSG ratio of plasma was highly significant already after 1 h of preparation delay (paired Student’s t-test p-value = 1.98 × 10^−5^, Fig. 1A, left panel). The same was true for GSH and GSSG content, with GSH being far more significant in this regard (paired Student’s t-test p-value (0 *versus* 60 min) for GSH and GSSG: 1.62 × 10^−12^ and 1.23 × 10^−5^, respectively).

**Figure 1.**
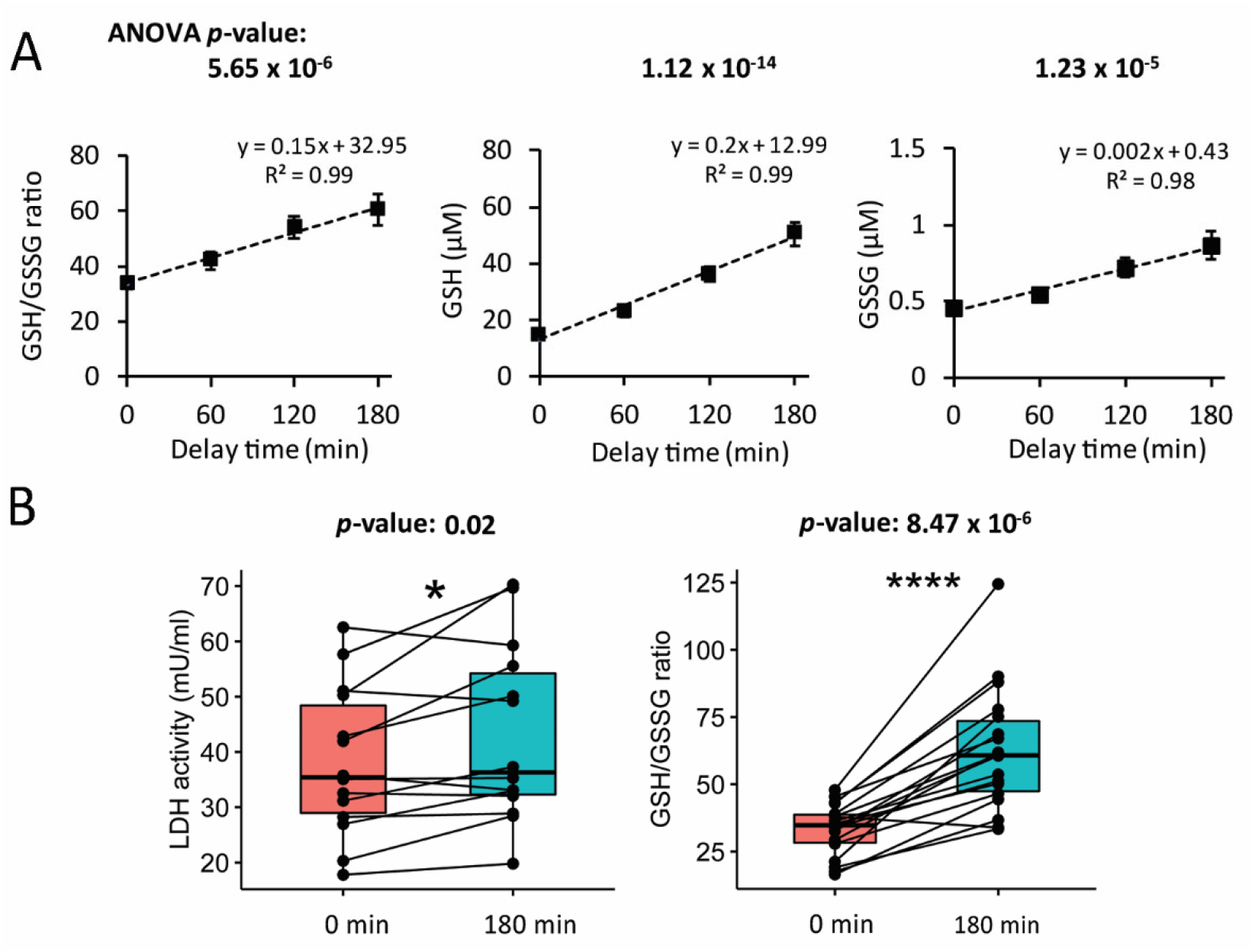
Concentrations of both GSH and GSSG increase over preparation delay time in plasma most likely due to hemolysis. A: GSH/GSSG ratio (left), absolute GSH (middle) and GSSG (right) content in plasma of 20 volunteers after their blood was incubated for 0, 60, 120 and 180 min prior plasma preparation. Points represent mean values (n = 19 or 20) ± S.E.M on which linear regression was applied. B: Lactate dehydrogenase activity assay (left, n = 14) suggest an increase in LDH activity in plasma over time (after 3 h of incubation on RT).

However, compared to the commonly used LDH activity assay, the GSH/GSSG ratio was a much more potent indicator of pre-analytical plasma preparation delay (right).

### Increase in glutathione was accompanied with higher lactate dehydrogenase activity of the plasma

We hypothesized that the observed increase in glutathione content after the delays could indicate increased cellular damage. To test this hypothesis, we carried out a lactate dehydrogenase (LDH) assay, a common test for cells undergoing necrosis, apoptosis and disruption of cellular membranes, resulting in the release of LDH (*18*). Thus, in our setting, an increase in LDH in plasma could indicate leakage from blood cells. Comparing LDH activity of plasma prepared minutes after blood collection (0 min time point) to plasma collected after a delay of 3 h (180 min time point; in pairwise manner i.e. per volunteer), LDH activity of plasma was significantly higher (Student t-test *p*-value = 0.02) at the later time point (Fig. 1B, left panel). However, pairwise comparison of individual GSH/GSSG ratios to LDH activities reveals higher significance of the GSH/GSSG ratio (paired Student’s t-test p-value = 8.47 × 10^−6^), indicating far better sensitivity of the GSH/GSSG ratio as a marker of prolonged blood incubation time prior plasma preparation (Fig. 1B, right panel).

### Common blood parameters were not affected by NEM spiked tubes

As glutathione needs to be protected from artificial oxidation immediately upon sample collection (*9, 10*), we next tested if addition of NEM directly to the blood tubes prior blood collection would interfere with common downstream clinical applications. For that purpose, we compared clinical routine parameters covering hematology, electrolytes and a number of biochemical compounds from blood collected to either NEM containing (2.5 mM final concentration) or NEM-free (control; PBS used as a vehicle) Li-heparin blood collection tubes (Fig. 2&3, Table 2 and Suppl. Data 1).

**Figure 2.**
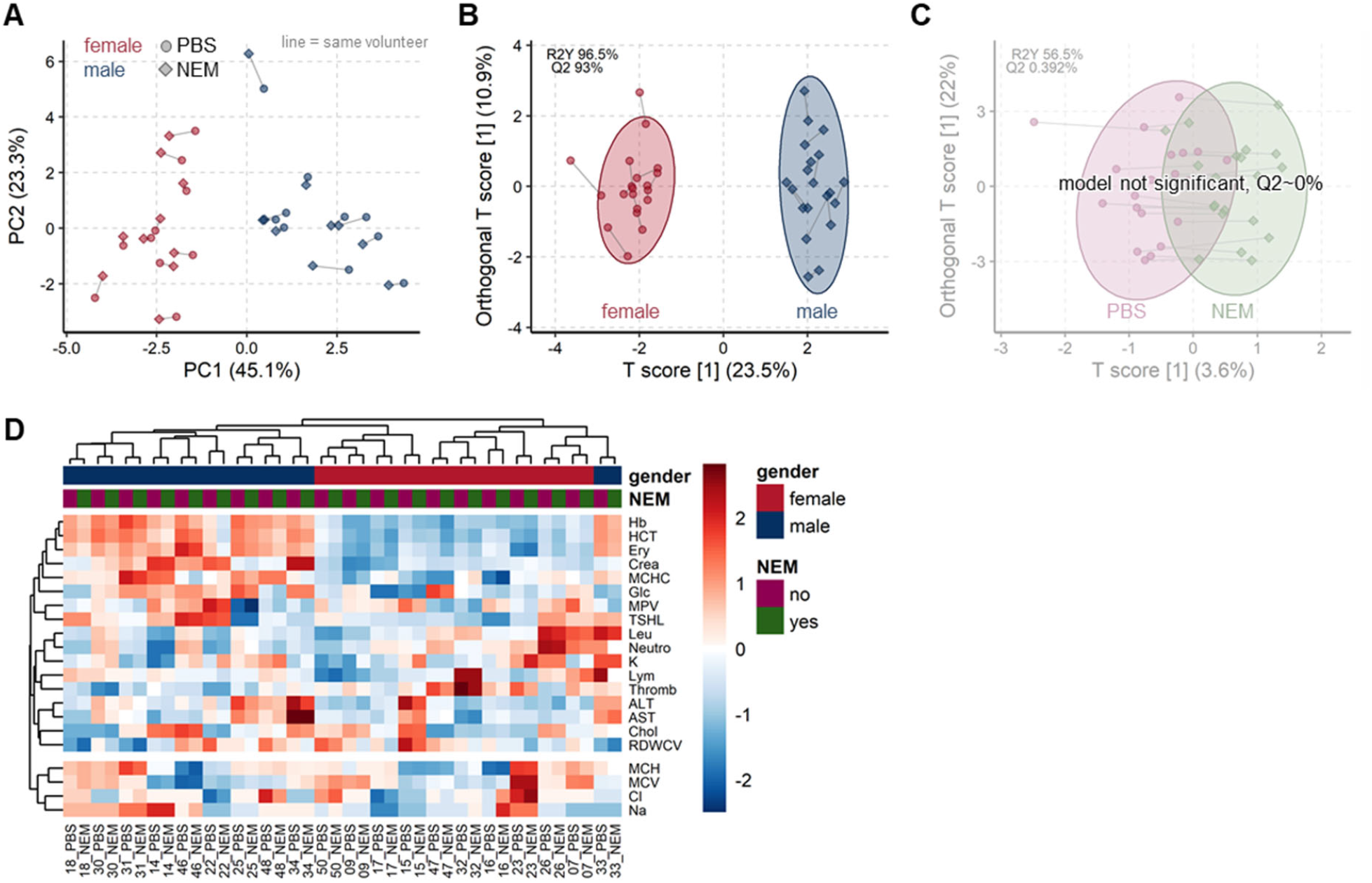
Addition of NEM to the blood tubes does not influence common clinical blood parameters. A: The unsupervised PCA scores plot shows a very high similarity of all 21 clinical parameters for each of the 20 volunteers (connected by grey line) when measured from tubes spiked with PBS (circles) or with NEM (diamonds). Samples are colored according to gender (female red, male blue) showing the expected strong group separation. B: OPLS-DA scores plot reconfirm the PCA findings, that clinical parameters significantly differ between gender, while C: the addition of NEM to PBS has no significant impact (A-C values in Suppl. Data 1). D: Heatmaps with hierarchical clustering show the similarity of obtained values between tubes with PBS or NEM, always clustering both samples from the same volunteer together.

**Figure 3.**
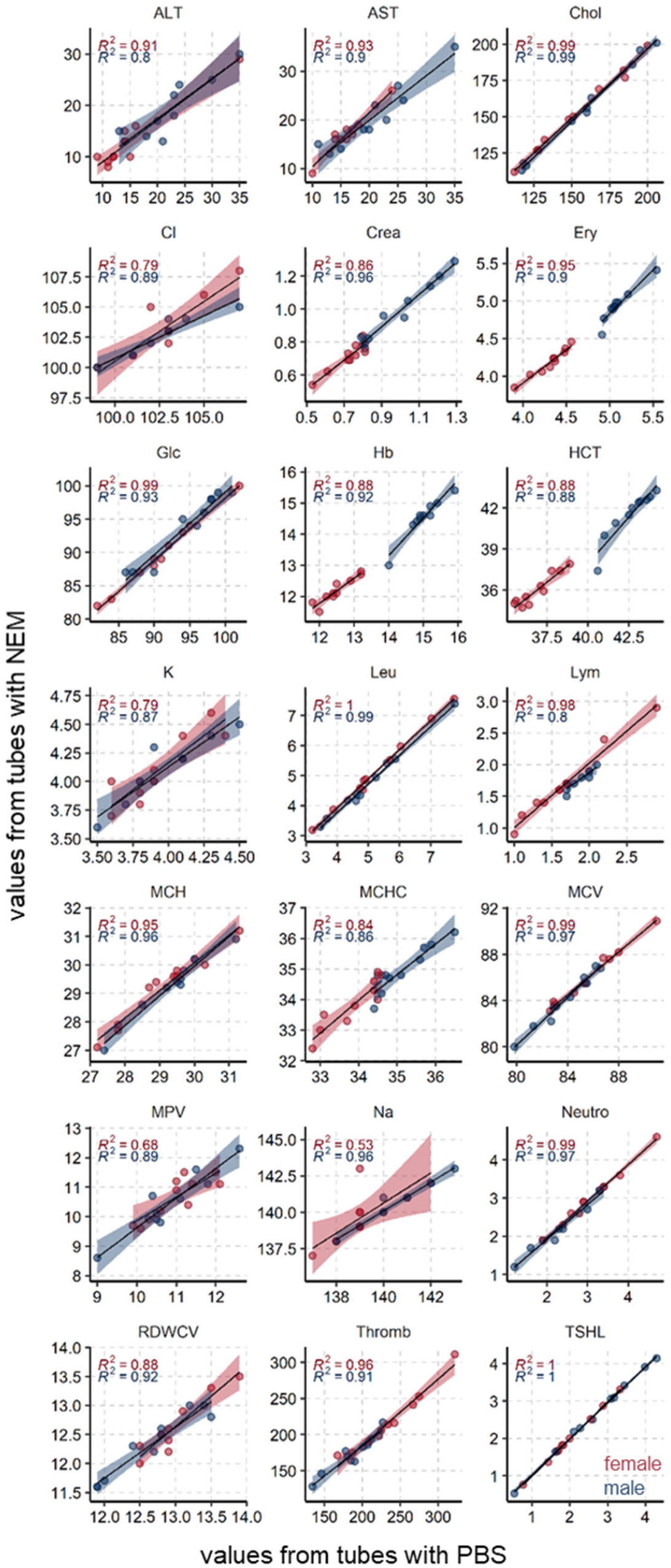
Direct comparison of clinical parameters measured from NEM or PBS spiked tubes. The linear correlation was calculated separately per gender, R^2^ are given in blue for males and red for females, the corresponding transparent stripes mark the 95% confidence intervals.

Comparison of the cell counts Baso, Eos, and Mono (for abbreviations see Table 2) were only qualitatively possible due to the categoric nature of the data. Most results were equal, if differences were found, these were within the expected measurement precision. The disease markers CRP and TNTHS were as expected low confirming the excellent health status of the included volunteers. The values were often below the limit of detection leading to too many missing values for statistical analysis. Thus, from the 26 assessed clinical parameters a total of 21 were available for statistical analysis.

We first employed the unsupervised multivariate method MacroPCA (*16*) to gain unbiased insight into the main driving factors. The scores plot shows that samples with NEM or PBS from the same volunteer were very near, hence very similar, to each other (Fig. 2A). Biological differences between females and males (factor gender) allow here to function as a positive control. The gender differences are well visible in the scores plot by a strong group separation along the first component (x-axis) which reflects 45% of the variability in the dataset. Hierarchical clustering confirmed the dominant similarity between tubes from the same volunteer by always clustering these beside each other (Fig. 2D). Overall samples were almost perfectly clustered by gender, emphasizing that gender differences are much larger than minor deviations between tubes from the same volunteer. OPLS-DA confirmed that gender differences in clinical parameters are highly significant (Fig. 2B) with a Q2 of 93 % far above the cut-off Q2 > 50 %, while no significant differences were found between PBS or NEM spiked tubes (Fig. 2C) with a Q2 of around 0 %.

In addition, we performed a direct comparison by linear correlation. Values from NEM and PBS tubes correlated in general very good with each other for both genders (Fig. 3). As expected, males had in general much higher red blood cell related values (Ery, Hb, HCT, MCHC, MCV) and higher creatinine (crea) as a result from higher muscle mass.

Furthermore, the majority of the parameters displayed a low mean within-subject coefficient of variation (CV_w_) of below 5 % (Table 2). Among the parameters with a CV_w_ higher than 5 % were alanine and aspartate aminotransferase (ALT and AST, CV_w_=11.7 and 5.7 %, respectively), platelets (CV_w_=5.47 %) and troponin (CV_w_=17.6 %). A detailed list of individual values of all measured parameters can be found in Suppl. Data 1.

## DISCUSSION

Delay between blood collection and plasma preparation can introduce variability into consequent downstream analysis of the plasma metabolome (*3, 4*) as well as proteome (*5*). SOPs for plasma preparation suggest that blood centrifugation should be carried out within 4 h upon blood collection (*19*), a time period which was shown to be sufficient to shift plasma metabolomic signatures (*4*). Nevertheless, the majority of studies investigating potential markers of the pre-analytical delay focused mainly on molecules which increase in plasma after or at the indicated four hour mark (*2*), which, for many downstream analyses, might already be detrimental.

In contrast, we here report GSH and/or GSH/GSSG ratio as potential early markers of plasma preparation delay. GSH and/or GSH/GSSG ratio plasma values rose significantly already after 1 h of blood incubation at RT prior to centrifugation (0 min *versus* 60 min time point paired Student t-test p-value of 1.98 × 10^−5^ for GSH/GSSG ratio and 1.62 × 10^−12^ for GSH). Over the monitored period of 3 h, we observed that both GSH and GSSG content in plasma increased linearly (R^2^=0.99 and 0.98, respectively, Fig. 1A) with longer centrifugation delay. However, as GSH increased faster than GSSG, this also led to a significant increase of the GSH/GSSG ratio in plasma. Therefore, from the technical perspective, both GSH and GSSG as well as the GSH/GSSG ratio constitute reliable indicators of time delays at RT prior to centrifugation for plasma collection.

One potential reason for the observed increase in glutathione content in plasma could be time dependent blood cell damage. It is known that incubation of blood in EDTA tubes at RT for a period of 24 h leads to approximately 2 % hemolysis (*20*), which, given the drastically higher GSH content of blood cells compared to the cell-free blood fraction, could be more than enough to cause a noticeable rise of GSH content in plasma already within the first hours of blood collection. In support of this hypothesis, an LDH activity assay demonstrated that 3 h of blood incubation indeed resulted in increased cellular damage (Fig. 1B, left panel). However, the time dependent rise in LDH activity was significant only at the three-hour time-point when tested in pair-wise manner (Student t-test p-value 0 versus 180 min (LDH activity) = 0.02, Fig. 1B), rendering it a far less sensitive marker than GSH or the GSH/GSSG ratio (Student t-test p-value 0 versus 180 min (GSH/GSSG ratio) = 8.47 × 10^−6^, Fig. 1B).

The second goal of this study was the incorporation of cysteine-quenchers into clinical routine, which would also enable precise redox status measurements. Therefore, we investigated if spiking standard blood tubes with 2.5 mM NEM would show any influence on the most common routinely used blood parameters. The results from NEM spiked tubes were found to be very similar to PBS spiked tubes (as vehicle control) with several different statistical methods. The unsupervised methods MacroPCA and hierarchical clustering found a very high similarity of values from PBS and NEM tubes, while the expected large difference between males and females was as well detected (Fig. 2). The significance was accordingly confirmed with the supervised machine learning method OPLS-DA. When directly comparing values from NEM to PBS tubes a clear linear correlation existed with mostly high R^2^ >0.9 (Fig. 3). Lower R^2^ (0.58-0.8, e.g. in Na, Cl, K, MPV) can be rather attributed to the instrument’s precision and small range of values. The majority of the tested parameters (Table 2) showed very low paired coefficients of variation with most CV_w_ < 5 %. The few exceptions were: ALT (CV_w_=11.7 %), AST (CV_w_=5.7 %) and platelets (CV_w_=5.5 %), which showed all R^2^ > 0.8. All of the listed CV_w_ are in accordance with the previously published within-subject, biological variance range for each of the parameters. For example, according to the currently available databases on biological variation (BV) (version from 1999 (*21*) and 2014 update (*22*)), as well as more recent systematic review (*23*), reported CV_w_ range for ALT is 11.1 % – 58 %, and 3.0 % – 24 % (*22*) or 32.3 % for AST (*23*). Regarding the platelets, the study from 2018 by Coskun et *al*., reported a mean CV_w_ value of 7.2 % for platelets (*24*). Values in this study were obtained from 30 individuals and corroborated the proposed CV_w_ ranges from the BV databases (*21, 22*). Altogether our results find no influence of NEM on the tested routine blood parameters.

## CONCLUSION

As a result of this study we propose that GSH, GSSG or the ratio of GSH/GSSG could be used as highly sensitive potential biomarkers for a centrifugation delay prior to plasma preparation. The study workflow and the results are summarized in Figure 4. In addition, we provide information regarding the rate of hemolysis for blood samples kept for up to three hours at RT. Lastly, in an attempt to aid implementation of cysteine-quenchers such as NEM into standard clinical routine for precise downstream redox analyses, we demonstrate that addition of NEM to the blood tubes does not influence common routine clinical parameters. Overall, this study provides guidelines how to track and consider pre-analytical variability, all in order to improve the confidence in the obtained results.

**Figure 4.**
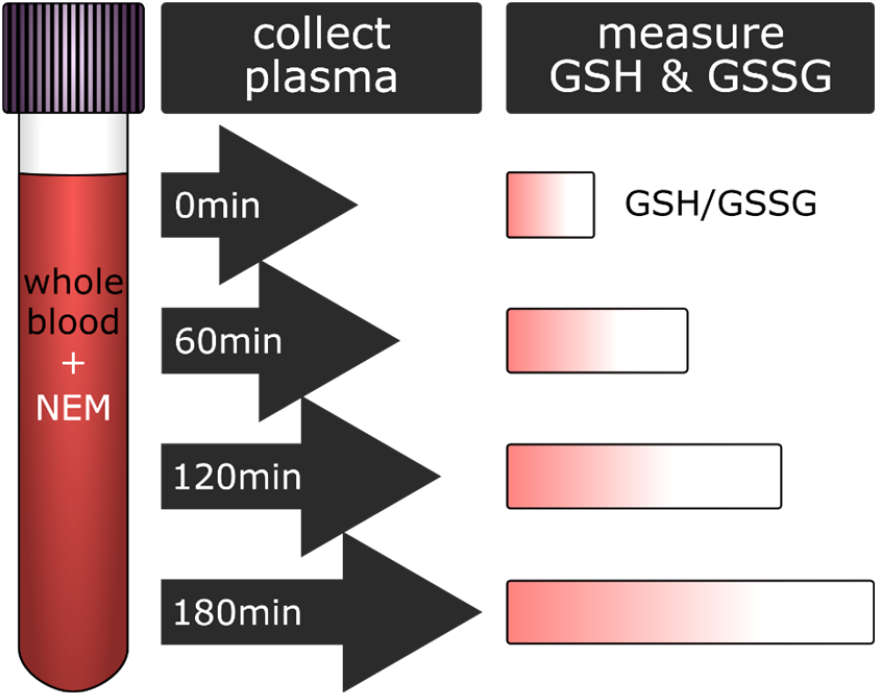
Summary of the workflow and the obtained results. Blood from 20 healthy volunteers was collected to pre-NEMylated blood collection tubes (2.5 mM final NEM concentration). At regular intervals (0, 60, 120 and 180 min) an aliquot of collected blood from each volunteer was spun down to collect the plasma, which was then used for further analysis and revealed an increase in GSH/GSSG ratio in a time dependent manner.

## Abbreviations

BMI: body mass index
BH: Benjamini-Hochberg
GSH: Glutathione
GSSG: glutathione disulfide
LDH: lactate dehydrogenase
MVA: multivariate analysis
NEM: N-ethylmaleimide
OPLS-DA: orthogonal projections to latent structures discriminant analysis
PCA: principal component analysis
RT: room temperature (20°C-25°C)
SD: standard deviation
SEM: standard error of mean
SOP: standard operating procedure
UVA: univariate analysis

## ACKNOWLEDGEMENT

We thank Rasa Vitonyte, Eva Svehlikova, Sigrid Deller, Elisabeth Langmann, Robert Stefan Lipp, Amar Alikadic, Jehona Qerimi-Hyseni, Ines Mursic, Andrea Halsegger, Michael Wolf, Martina Urschitz, Ana Semonik, Martina Brunner, Rene Peter Engel, Beatrix Stroisnik, Georg Wiesnegger, Sven Miedler, Annemarie Marold, Julia Matejka, Franziska Vogl, Martin Saxinger, Selina Kofler, Martina Tomberger, and Jessica Schweiger for their excellent support of the observational study. We thank Shimadzu Corporation, Kyoto, Japan for their excellent scientific cooperation.

The samples/data used for this project have been provided by Biobank Graz, Austria.

